# The carbon footprint of bioinformatics

**DOI:** 10.1101/2021.03.08.434372

**Authors:** Jason Grealey, Loïc Lannelongue, Woei-Yuh Saw, Jonathan Marten, Guillaume Meric, Sergio Ruiz-Carmona, Michael Inouye

**Affiliations:** Cambridge Baker Systems Genomics Initiative, Baker Heart and Diabetes Institute, Melbourne, Victoria, Australia; Department of Mathematics and Statistics, La Trobe University, Melbourne, Australia; Cambridge Baker Systems Genomics Initiative, Department of Public Health and Primary Care, University of Cambridge, Cambridge, UK; British Heart Foundation Cardiovascular Epidemiology Unit, Department of Public Health and Primary Care, University of Cambridge, Cambridge, UK; Health Data Research UK Cambridge, Wellcome Genome Campus and University of Cambridge, Cambridge, UK; Department of Infectious Diseases, Central Clinical School, Monash University, Melbourne Australia; British Heart Foundation Centre of Research Excellence, University of Cambridge, Cambridge, UK; National Institute for Health Research Cambridge Biomedical Research Centre, University of Cambridge and Cambridge University Hospitals, Cambridge, UK; The Alan Turing Institute, London, UK

**Keywords:** carbon footprint, bioinformatics, genomics, green algorithms

## Abstract

Bioinformatic research relies on large-scale computational infrastructures which have a non-zero carbon footprint. So far, no study has quantified the environmental costs of bioinformatic tools and commonly run analyses. In this study, we estimate the bioinformatic carbon footprint (in kilograms of CO_2_ equivalent units, kgCO_2_e) using the freely available Green Algorithms calculator (www.green-algorithms.org). We assess (i) bioinformatic approaches in genome-wide association studies (GWAS), RNA sequencing, genome assembly, metagenomics, phylogenetics and molecular simulations, as well as (ii) computation strategies, such as parallelisation, CPU (central processing unit) vs GPU (graphics processing unit), cloud vs. local computing infrastructure and geography. In particular, for GWAS, we found that biobank-scale analyses emitted substantial kgCO_2_e and simple software upgrades could make GWAS greener, e.g. upgrading from BOLT-LMM v1 to v2.3 reduced carbon footprint by 73%. Switching from the average data centre to a more efficient data centres can reduce carbon footprint by ~34%. Memory over-allocation can be a substantial contributor to an algorithm’s carbon footprint. The use of faster processors or greater parallelisation reduces run time but can lead to, sometimes substantially, greater carbon footprint. Finally, we provide guidance on how researchers can reduce power consumption and minimise kgCO_2_e. Overall, this work elucidates the carbon footprint of common analyses in bioinformatics and provides solutions which empower a move toward greener research.

## Introduction

Biological and biomedical research now requires the analysis of large and complex datasets, which wouldn’t be possible without the use of large-scale computational resources. Whilst bioinformatic research has enabled major advances in the understanding of a myriad of diseases such as cancer [1]–[3] and COVID-19 [4], the costs of the associated computing requirements are not limited to the financial; the energy usage of computers causes greenhouse gas (GHG) emissions which themself have a detrimental impact on human health.

Energy production affects both human and planetary health. The yearly electricity usage of data centres and high performance computing (HPC) facilities (200 TWh [5]) already exceeds the consumption of countries such as Ireland or Denmark [6] and is predicted to continue to rise over the next decade [5], [7]. Power generation, through the associated emissions of GHGs, is one of the main causes of both outdoor air pollution and climate change. Every year, it is estimated that 4.2 million deaths are caused by ambient air pollution alone while 91% of the world’s population suffers from air quality below the World Health Organisation standards [8]. Global warming results in further consequences on human health, economy and society: the daily population exposure to wildfires has increased in 77% of countries [9], 133.6 billion potential work hours were lost to high temperatures in 2018 and with 220 million heatwave exposures, vulnerable populations (aged 65 and older) are affected at an unprecedented level.

The growth of large biological databases, such as UK Biobank [10], All of Us Initiative [11], and Our Future Health [12], has substantially increased the need for computational resources to analyse these data and will continue to do so. With climate change an urgent global emergency, it is important to assess the carbon footprint of these analyses and their requisite computational tools so that environmental impacts can be minimised.

In this study, we estimate the carbon footprint of common bioinformatic tools using a model which accounts for the energy use of different hardware components and the emissions associated with electricity production. For each analysis, we contextualise the carbon footprint in multiple ways, such as distances travelled by car or with regards to carbon sequestration by trees. This study raises awareness, provides easy-to-use metrics, and makes recommendations for greener bioinformatics.

## Results

We estimated the carbon footprint of a variety of bioinformatic tools and analyses (**Table 1, Table 2**) using the Green Algorithms model and online tool (**Methods**). For each software, we utilised benchmarks of running time and computational resources; in the rare cases where published benchmarks were unavailable, we used in-house analyses to estimate resource usage (**Methods**). The estimations are based on the global average data centre efficiency (PUE) of 1.67 [13], the global average carbon intensity (0.475 kgCO_2_e/kWh [14]) and a usage factor of 1 (**Methods**).

**Table 1:**
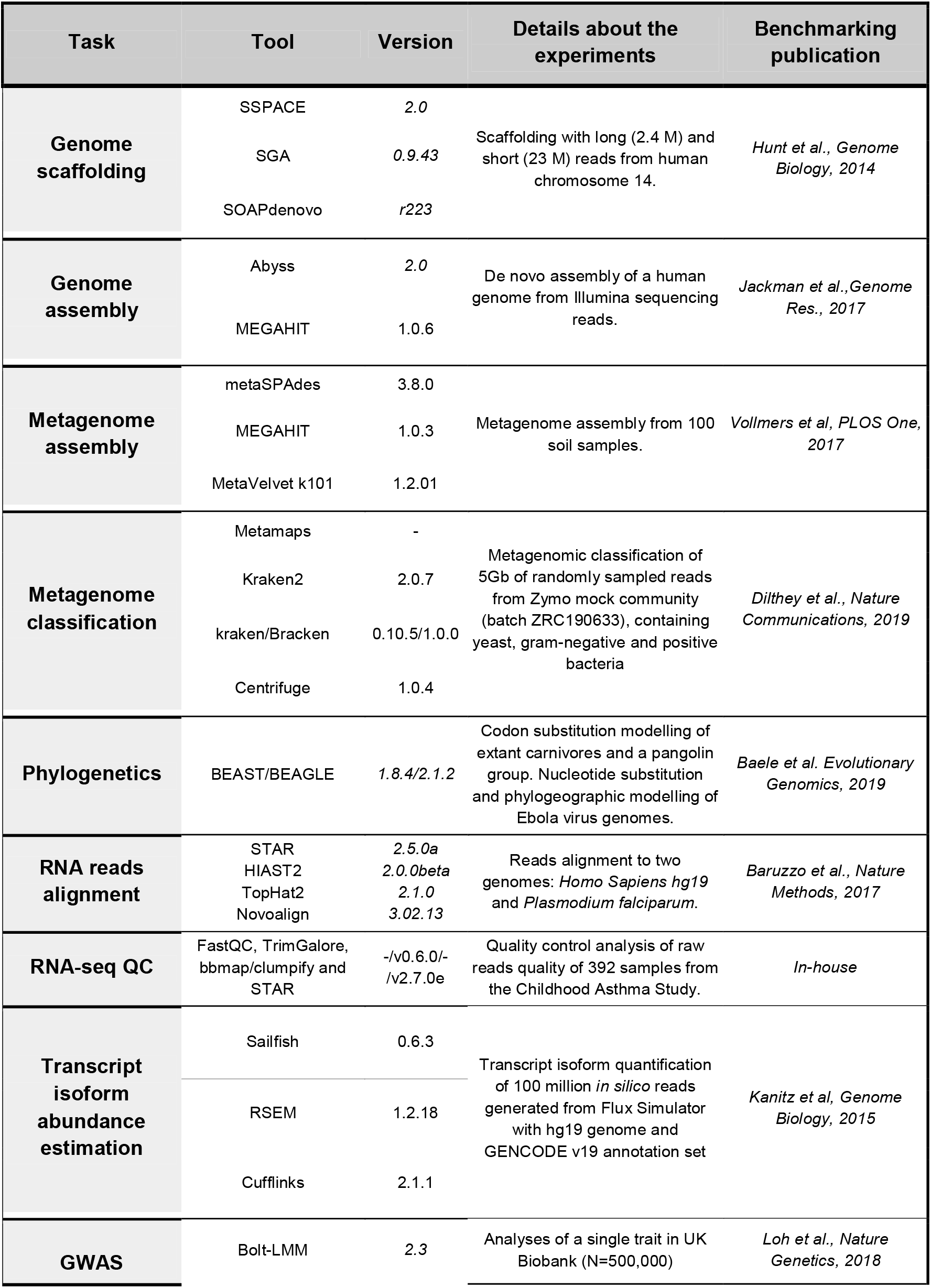
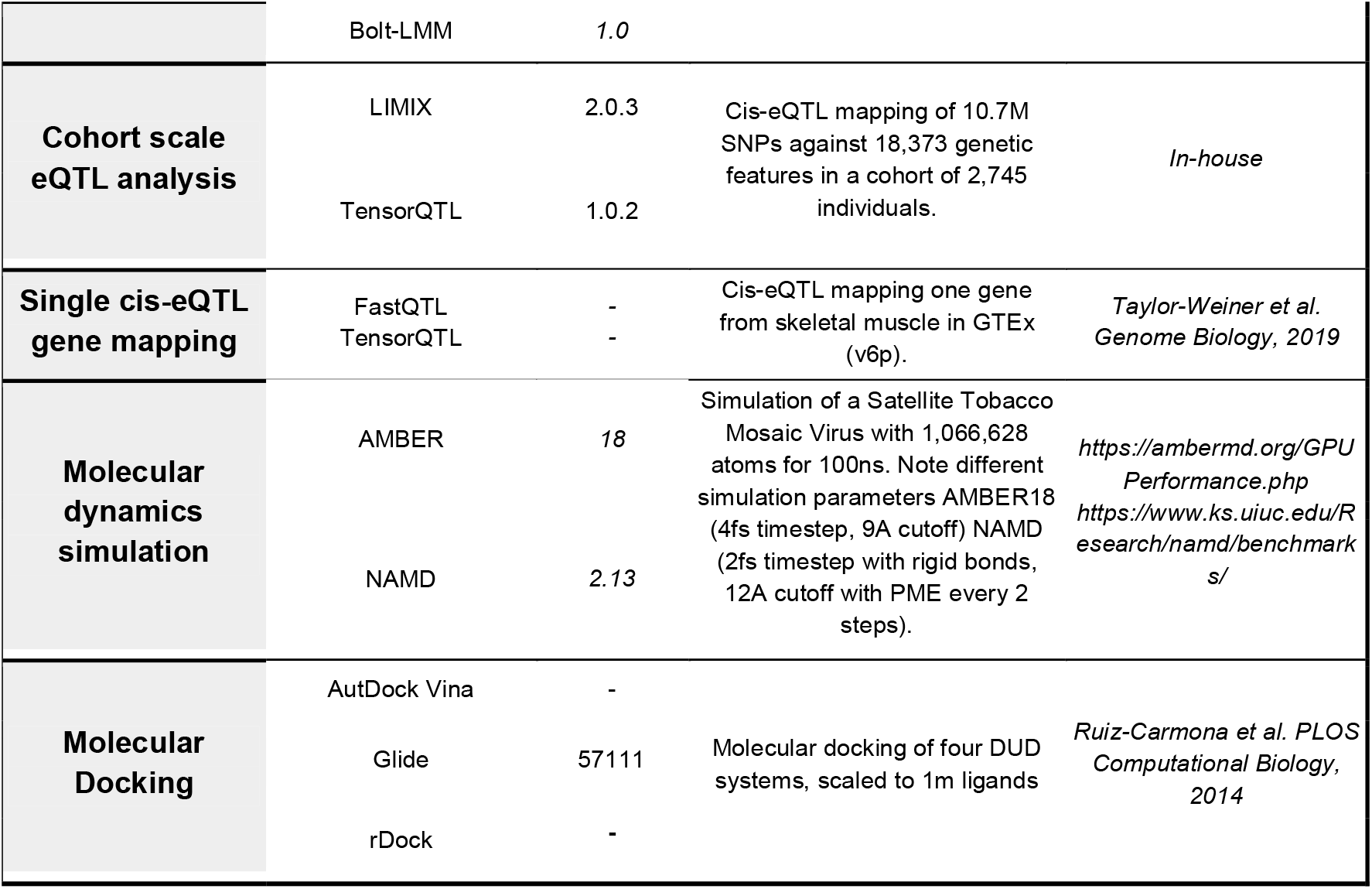
A description of the tasks, tools and experiments used in this study.

**Table 2:** The estimated carbon footprint of bioinformatic tasks. This table details and contextualises the carbon footprint of the tasks detailed in Table 1. In addition to the carbon footprints are the number of tree-months it would take an adult tree to sequester the CO_2_, and the number of kilometres one could travel in an average European car to output the same amount of CO_2_. *These methods were estimated in-house and not from a published benchmark.

| Task | Tool | Carbon footprint (kgCO<sub>2</sub>e) | tree-months | km in a car (EU) |
| --- | --- | --- | --- | --- |
| Genome scaffolding (long read) | SGA | 0.0293 | 0.0319 | 0.2 |
|  | SSPACE | 0.0010 | 0.0011 | 0.01 |
|  | SOAPdenovo2 | 0.0015 | 0.0016 | 0.01 |
| Genome scaffolding (short read) | SGA | 0.1302 | 0.1420 | 0.7 |
|  | SSPACE | 0.0027 | 0.0029 | 0.02 |
|  | SOAPdenovo2 | 0.0036 | 0.0039 | 0.02 |
| De novo assembly of one human genome | Abyss2.0 | 10.66 | 11.63 | 60.9 |
|  | MEGAHIT | 15.11 | 16.48 | 86.3 |
| Metagenome assembly | metaSPAdes | 186.46 | 203.41 | 1,065.5 |
|  | MEGAHIT | 76.81 | 83.79 | 438.9 |
|  | Meta Velvet k101 | 14.28 | 15.58 | 81.6 |
| Metagenome classification (short read) | Centrifuge | 0.013 | 0.0138 | 0.1 |
|  | Kraken2 | 0.0052 | 0.0057 | 0.03 |
|  | Kraken/Bracken | 0.092 | 0.1000 | 0.5 |
| Metagenome classification (long read) | MetaMaps | 18.25 | 19.91 | 104.3 |
| RNA read alignment *Homo Sapiens hg19* | STAR v2.5.0a | 0.0097 | 0.0105 | 0.1 |
|  | HISAT2 | 0.0054 | 0.0059 | 0.03 |
|  | TopHat2 | 0.3173 | 0.3461 | 1.8 |
|  | Novoalign | 0.9766 | 1.0653 | 5.6 |
| RNA read alignment *Plasmodium falciparum* | STAR v2.5.0a | 0.3693 | 0.4029 | 2.1 |
|  | HISAT2 | 0.0052 | 0.0057 | 0.03 |
|  | TopHat2 | 0.2394 | 0.2612 | 1.4 |
|  | Novoalign | 0.6710 | 0.7320 | 3.8 |
| *RNA sequencing quality control pipeline | FastQC + TrimGalore + clumpify + STARv2.7.0e | 54.97 | 59.97 | 314.1 |
| Transcript isoform abundance estimation | Cufflinks - 1 core | 0.045 | 0.049 | 0.3 |
|  | RSEM - 1 core | 0.57 | 0.63 | 3.3 |
|  | Sailfish - 1 core | 0.0081 | 0.0088 | 0.05 |
|  | Cufflinks - 16 cores | 0.27 | 0.30 | 1.6 |
|  | RSEM - 16 cores | 1.40 | 1.53 | 8.0 |
|  | *Sailfish - 16 core* | 0.036 | 0.039 | 0.2 |
| **GWAS on a biobank with 1 trait** | *Bolt-LMM v1* | 17.29 | 18.86 | 98.8 |
|  | *Bolt-LMM v2.3* | 4.70 | 5.13 | 26.9 |
| **\*eQTL mapping for a cohort** | *TensorQTL* | 2.04 | 2.22 | 11.6 |
|  | *LIMIX* | 190.73 | 208.07 | 1,089.9 |
| **cis-eQTL mapping for 1 gene** | *FastQTL* | 0.0002 | 0.0002 | 0.001 |
|  | *TensorQTL* | 0.00001 | 0.00001 | 0.00004 |
| **Virus molecular dynamics simulations** | *AMBER18* | 17.85 | 19.47 | 102.0 |
|  | *NAMD 2.13* | 95.19 | 103.84 | 543.9 |
| **Molecular docking** | *AutoDock Vina* | 514.12 | 560.86 | 2,937.9 |
|  | *Glide* | 12.90 | 14.07 | 73.7 |
|  | *rDock* | 153.71 | 167.69 | 878.4 |

We considered a wide range of bioinformatic analyses: genome assembly, metagenomics, phylogenetics, RNA sequencing, genome-wide association analysis, molecular simulations and virtual screening. Detailed results are provided for each analysis below. Furthermore, we show that choices of hardware and software versions substantially affect the carbon footprint of a given analysis, in particular cloud vs. local computing platforms, memory usage, processor options, and parallel computing. These results provide, for each task, reference values of carbon footprints for researchers; however, we note how the estimations are likely to scale with different parameters (e.g. sample size or number of features) and ultimately would advise researchers to utilise the GA tool (www.green-algorithms.org).

### Genome assembly

Genome assembly is the process by which sequencing reads (short or long reads, or a combination) are combined to arrive at a single or set consensus sequences for an organism. Hunt et al. [15] compared SSPACE [16], SGA [17] and SOAPdenovo2 [18] for genome scaffolding using contigs produced with the Velvet assembler [19] and the human chromosome 14 GAGE dataset [20]; two read sets were compared, one using 22.7 million short reads (fragment length of 3 kb) and the other 2.4 million long reads (35 kb). Scaffolding the short reads resulted in 0.13, 0.0036, and 0.0027 kgCO_2_e when using SGA, SOAPdenovo2 and SSPACE, respectively (**Table 2**), which is equivalent to 0.14, 0.0039 and 0.0029 tree-months. For long reads scaffolding, the corresponding carbon footprints were lower, 0.029, 0.0015 and 0.0010 kgCO_2_e (0.032 to 0.0011 tree-months). As the running time of a number of genome assembly tools scale linearly with the number of reads [21], these results equate to between 0.0001 to 0.006 kgCO_2_e (0.0001 to 0.006 tree-months) per million short reads assembled and 0.0004 to 0.0122 kgCO_2_e (0.0005 to 0.0133 tree-months) per million long reads assembled. On average, long read assembly had a carbon footprint 3.2x larger than short-read assembly for the tools we measured. All three methods had similar performance on these read sets with SOAPdenovo2 slightly outperforming SGA and SSPACE.

For whole genome assembly of humans, the well-established softwares Abyss [22] and MEGAHIT [23] were benchmarked by Jackman et al. [22] using Illumina short read sequencing (815M reads, 379M uniquely mapped reads, 6kbp mean insert size) (**Table 2**). We estimated that this task emits 10.7 kgCO_2_e using Abyss and 15.1 kgCO_2_e using MEGAHIT (equivalent to 12 and 16 tree-months) and per million reads, 0.013 kgCO_2_e (Abyss2.0, 0.014 tree-months) and 0.019 kgCO_2_e (MEGAHIT, 0.020 tree-months).

### Metagenomics

Metagenomics is the sequencing and analysis of all genetic material in a sample. Based on a benchmark from Vollmers et al. [24], we estimated the carbon footprint of metagenome assembly with three commonly used assemblers, metaSPAdes [25], MEGAHIT [23] and MetaVelvet (k-mer length 101bp) [26] on 100 samples from forest soil (33M reads, median length 360 bp). We found carbon footprints ranged between 14 and 186 kgCO_2_e (16 and 203 tree-months), corresponding to 0.14 to 1.9 kgCO_2_e (0.2 to 2 tree-months) per sample. Meta-SPAdes had the greatest carbon footprint but also the best performance followed by MetaVelvet and MEGAHIT, respectively (**Table 2**).

For metagenomic classifiers, Dilthey et al. [27] benchmarked MetaMaps [27], Kraken2 [28], Kraken/Bracken [29], [30], and Centrifuge [31]. They compared these tools on ~5Gb of randomly sampled reads from an Oxford Nanopore GridION sequencing run from Zymo mock communities, which comprises five Gram-positive bacteria, three Gram-negative bacteria and two types of yeast. Carbon footprints differed by several orders of magnitude, MetaMaps had the largest footprint with 18.25 kgCO_2_e (19.9 tree-months), followed by Kraken/Bracken 0.092 kgCO_2_e (0.1 tree-months), Centrifuge 0.013 kgCO_2_e (0.014 tree-months) and Kraken2 0.0052 kgCO_2_e (0.0057 tree-months) (**Table 2**). The carbon footprints of metagenomic classification ranged from 0.001 to 0.018 kgCO_2_e (0.001 to 0.02 tree-months) per Gb of classified reads using short read classifiers (Kraken2, Centrifuge, Kraken/Bracken). Kraken2 had the highest performance over all taxonomic ranks when all reads were assembled, followed by Kraken/Bracken, Centrifuge and MetaMaps. However, when considering reads >1000bp, MetaMaps had the highest precision and recall for all available taxonomic levels, followed by Kraken2, Kraken/Bracken, and Centrifuge.

### Phylogenetics

Phylogenetics is the use of genetic information to analyse the evolutionary history and relationships amongst individuals or groups. Baele et al. [32] benchmarked nucleotide-based phylogenetic analyses with and without spatial location information to study the evolution of the Ebola virus during the 2013-2016 West African epidemics (1,610 genomes, 18,992 nucleotides [33]). The authors also investigated more complex codon models. For all these tasks, they utilised BEAST combined with BEAGLE [34].

We estimated the carbon footprint of nucleotide-based modelling of the Ebola virus dataset was between 0.01 to 0.08 kgCO_2_e depending on hardware choices (0.013 to 0.083 tree-months) without modelling spatial information and 0.07 to 0.3 kgCO_2_e (0.077 to 0.33 tree-months) when including it. More complex codon modelling of extant carnivores and pangolins resulted in a greater footprint, from 0.02 to 0.1 kgCO_2_e (0.02 to 0.1 tree-months) (**Figure 2, Supplementary table 2**). These results illustrate a trade-off between running time and carbon footprints, and we discuss it in more detail below (**Parallelisation, Processors**). It should be noted that the running time of BEAST, and therefore its carbon footprint, scales as a power law, that is, non-linearly with the number of loci [35].

### RNA sequencing

RNA sequencing (RNAseq) is the sequencing and analysis of all RNA in a sample. We first assessed the read alignment step in RNAseq using an extensive benchmarking by Baruzzo et al. [36]. We estimated the carbon footprint of aligning 10 million simulated 100-base read pairs to two different genomes, *Homo Sapiens* (hg19) and *Plasmodium falciparum* [36], which have substantially differing levels of complexity (*P. falciparum* with higher rates of polymorphisms and errors). The three most-cited software tested, STAR [37], HISAT2 [38] and TopHat2 [39], all had low recall on the malaria dataset, so we also assessed Novoalign [40] as it performed significantly better for this task (**Table 2**). Despite its greater performance for *P. falciparum*, Novoalign had the highest carbon footprint (0.67 kgCO_2_e, 0.73 tree-months) followed by STAR (0.37 kgCO_2_e, 0.40 tree-months), TopHat2 (0.24 kgCO_2_e, 0.26 tree-months) and HISAT2 with the lowest (0.0052 kgCO_2_e, 0.0057 tree-months). For human read alignment, all four methods had high recall. HISAT2 had, again, the lowest carbon footprint with 0.0054 kgCO_2_e (0.0059 tree-months) followed by STAR with 0.0097 kgCO_2_e (0.011 tree-months), TopHat2 with 0.32 kgCO_2_e (0.35 tree-months) and Novoalign with 0.98 kgCO_2_e (1.1 tree-months). As alignment tools are often reported with alignment speed (reads aligned in a given time) [37], [38], the carbon footprints of the analyses above scale accordingly and ranged from 0.001 to 0.1 kgCO_2_e (0.001 to 0.1 tree-months) per million human or *P. falciparum* reads.

To quantify the carbon footprint of a full quality control pipeline with FastQC, we utilised 392 RNAseq read sets obtained from PBMC samples [41], [42], with a median depth of 45 million paired-end reads and average length 146bp. Adapters were trimmed with TrimGalore [43], followed by the removal of optical duplicates using bbmap/clumpify [44]. Reads were then aligned to the human genome reference (Ensemble GRCh 38.98) using STAR [37]. We estimated the carbon footprint of this pipeline to be 55 kgCO_2_e (60 tree-months) for the full dataset, or 1.2 kgCO_2_e (1.3 tree-months) per million reads (**Table 2**), which scales linearly (**Additional file 2**).

For transcript isoform abundance estimation, we could assess Sailfish [45], RSEM [46], and Cufflinks [47] using the benchmark from Kanitz et al. [48] on simulated human RNA-seq data (hg19). The Flux Simulator software [49] and GENCODE [50] were used to generate 100 million single-end 50bp reads. The carbon footprints of this task were between 0.0081 and 1.4 kgCO_2_e (0.009 to 1.5 tree-months). Sailfish had the lowest footprint, followed by Cufflinks and RSEM. (**Table 2**). Kanitz et al. showed that the time complexity for most of the tools tested was approximately linear, i.e. the carbon footprint is proportional to the number of reads. Additionally, these tools offer the option of parallelisation. However, for example, the decrease in running time when using 16 cores instead of one was not sufficient to offset the increase in power consumption, which resulted in a 2- to 6-fold increase in carbon footprint when utilising 16 cores (**Table 2**). RSEM and Sailfish had similar performance in this benchmark, but Sailfish’s carbon footprint was 71-fold less than RSEM’s when using 1core and 39-fold less with 16 cores. This difference in carbon footprint was partly due to Sailfish not performing a read alignment step. Lastly, whilst Cufflinks is largely used for abundance estimation, its main purpose is transcript isoform assembly, resulting in a significantly lower accuracy here (at a higher carbon cost).

### Genome-wide association analysis

Genome-wide association analysis aims to identify genetic variants across the genome associated with a phenotype(s). Here, we assessed both genome-wide association studies (GWAS) and expression qualitative trait locus (eQTL) mapping in *cis*. We estimated the carbon footprint of GWAS with two different versions of Bolt-LMM [51] on the UK Biobank [10] (500k individuals, 93M imputed SNPs). We found that a single trait GWAS would emit 17.3 kgCO_2_e (18.9 tree-months) with Bolt-LMM v1 and 4.7 kgCO_2_e (5.1 tree-months) with Bolt-LMM v2.3 (**Table 2**), a reduction of 73%. GWAS typically assess multiple phenotypes, e.g. metabolomics GWAS consider several hundred to thousands of metabolites; since the association models in GWAS are typically fit on a per-trait basis, the carbon footprint is proportional to the number of traits analysed. Bolt-LMM’s carbon footprint also scales linearly with the number of genetic variants [52], meaning that biobank-scale GWAS using UK Biobank (500k individuals) has a carbon footprint of 0.05 kgCO_2_e per million variants (0.06 tree-months) with Bolt-LMM v2.3 and 0.2 kgCO_2_e per million variants (0.2 tree-months) with Bolt-LMM v1. However, Bolt-LMM doesn’t scale linearly with the number of samples (*time ~ O(N^1.5^)* [52]), which must be taken into account when scaling the values to a different sample size.

For cis-eQTL mapping, we compared the carbon footprint using either CPUs or GPUs on two example datasets, first on a small scale using skeletal muscle data from GTEx [53] (1 gene, 700 individuals) with a benchmark of FastQTL (CPU) [54] and TensorQTL (GPU) [55], [56] from Taylor-Weiner et al. [56]. Secondly, we used an in-house assessment (**Methods**), to estimate the carbon footprint of a CPU-based analysis with LIMIX [57] to GPU-based TensorQTL using a larger cohort of 2,745 individuals with 18k genetic features and 10.7m SNPs (**Table 2**). In both cases, footprints were lower using GPUs instead of CPUs. The carbon footprint for the smaller scale GTEx benchmark was 28 times smaller when utilising the GPU instead of the CPU method: 0.0002 kgCO_2_e (0.0002 tree-months) with FastQTL, 0.00001 kgCO_2_e (0.00001 tree-months) with TensorQTL. Similarly, for the cohort scale cis-eQTL mapping, the carbon footprints were 94 times smaller when utilising the GPU approach: 191 kgCO_2_e (208 tree-months) with LIMIX and 2 kgCO_2_e (2 tree-months) with TensorQTL. The scaling of eQTLs is complex, and the carbon footprint doesn’t scale linearly with the number of traits or sample size [56], [57].

### Molecular simulations and virtual screening

Molecular simulations and virtual screening are the use of computational simulation to model and understand molecular behaviour and the *in silico* scanning of small molecules for the purposes of drug discovery. We estimated the carbon footprint of simulating molecular dynamics with the Satellite Tobacco Mosaic Virus (1,066,628 atoms) for 100ns [58], [59] to be 17.8 kgCO_2_e (19 tree-months) using AMBER [60] and 95 kgCO_2_e (104 tree-months) using NAMD [61] (**Table 2**). This corresponds to 1 kgCO_2_e per ns (1 tree-month) when utilising NAMD and 0.2 kgCO_2_e per ns (0.2 tree-months) with AMBER. There are small discrepancies between the simulation parameters used by the tools (**Table 1**) so they can’t be compared directly. Furthermore, due to a lack of information, neither of these estimations include the power usage from memory.

Using a benchmark from Ruiz-Carmona et al. [62], we estimated the carbon footprint of three molecular docking methods, AutoDock Vina, Glide and rDock [62]–[64]. The data are based on the directory of useful decoys (DUD) benchmark set [65]. This study tested the three docking methods on four DUD systems ADA, COMT, PARP, and Trypsin. Where we used the average computational running time on these four DUD systems to estimate the carbon footprint of a 1 million ligand campaign. Glide, the fastest but not freely available tool had the smallest carbon footprint with 13 kgCO_2_e (14 tree-months), whilst rDock, which is freely available, had a footprint of 154 kgCO_2_e (168 tree-months), and AutoDock Vina (also freely available) had the largest impact with 514 kgCO_2_e (561 tree-months) (**Table 2**). rDock was the lowest carbon emitting method that was freely available and had comparable performance to Glide [62].

### Efficiency of local data centres, geography and cloud computing

Cloud computing facilities and large data centres are optimised to significantly reduce overhead power consumption such as cooling and lighting. A report from 2016 estimated that energy usage by data centres in the US could be reduced by 25% if 80% of the smaller data centres were aggregated into larger and more efficient data centres (hyperscale facilities) [66]. This was consistent with the distribution of PUEs (**Methods**): compared to the global average PUE of 1.67, Google Cloud’s PUE of 1.11 [67] reduces the carbon footprint of a task by 34%. Other cloud providers also achieve low PUEs, Microsoft Azure reduces the carbon footprint by 33% (PUE=1.125 [68]) and Amazon Web Service by 28% (PUE=1.2 [69]).

The use of cloud facilities may also enable further reductions of carbon footprint by allowing for choice of a geographic location with relatively low carbon intensity. While the kgCO_2_e for specific analyses utilising cloud or local data centre platforms are best estimated with the Green Algorithm calculator (www.green-algorithms.org), we found that a typical GWAS of UK Biobank considering 100 traits using the aforementioned GWAS framework (see **Genome-wide association analysis**) together with BoltLMM v2.3 on a Google Cloud server in the UK would lower the carbon footprint by 81% when compared to the average local data centre in Australia (**Figure 1**), potentially saving 705 kgCO_2_e (769 tree-months).

**Figure 1.**
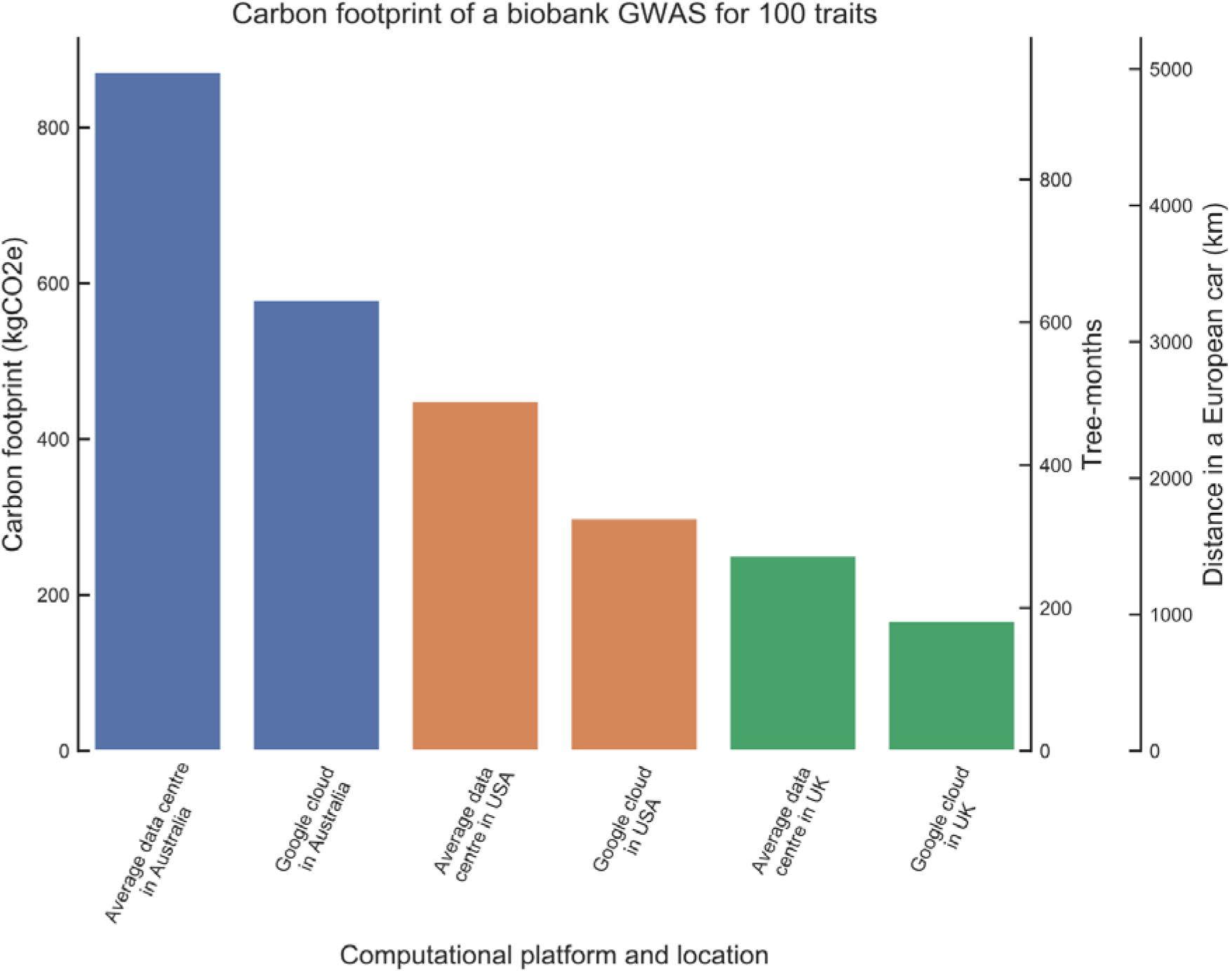
Location and computational platforms affect carbon footprint. This plot details the carbon footprint (in kgCO_2_e, tree-months, and European car km) of a biobank scale 100 trait GWAS in various locations and platforms. Average data centres have a PUE of 1.67 [13], Google cloud has PUE of 1.11[67], Australia has a carbon intensity of 0.88 kgCO_2_e/kWh, USA 0.453 kgCO_2_e/kWh, and UK 0.253 kgCO_2_e/kWh [74].

### Parallelisation

Numerous algorithms use parallelisation to share the workload between several computing cores and reduce the total running time. However, this can increase carbon footprint [70] and we found that parallelisation frequently results in tradeoffs between running time and carbon footprint. In some cases, the reduction in running time is substantial. For example, executing the phylogenetic codon model (**Phylogenetics**) on a single core would take 7.8 hours and emit 0.066 kgCO_2_e, but with two cores, the carbon footprint increased by 4% while running time was decreased by 46% (1.9x speedup). With 12 cores, run time decreased 86% (7.2x speedup) but the carbon footprint increased by 57%. In other cases, speedup was marginal, e.g. the phylogeographic model had a running time of 3.86 hours with a carbon footprint of 0.070 kgCO_2_e when using two cores (**Figure 2**). Increasing the parallelisation to 10 cores reduced run time by only 5% but increased carbon footprint by 4-fold.

**Figure 2:**
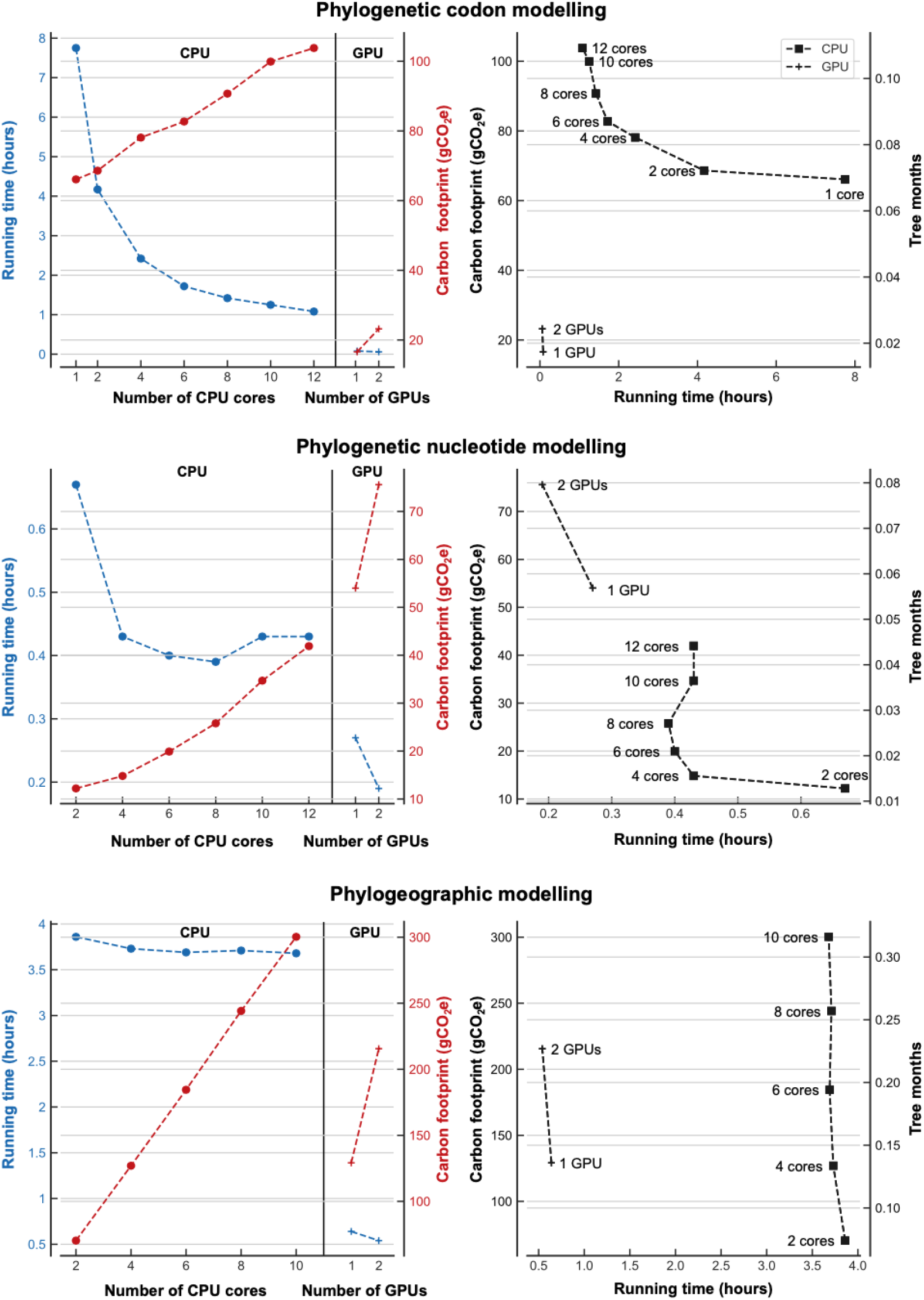
The effect of hardware choices and parallelisation on carbon footprint. The carbon footprint of BEAST/Beagle implemented on multi-core CPU or GPUs for three different tasks. The plots on the left detail both the running time and carbon footprint against the number of cores utilised. The plots on the right detail the running time solely against carbon footprint (contextualised with tree-months) for both CPUs and GPUs. The numerical data is available in **Supplementary Table 2**.

### Memory

Memory’s power consumption depends mainly on the memory available, not on the memory used [70], [71]; thus, having too much memory available for a task results in unnecessary energy usage and GHG emissions. Although memory is usually a fixed parameter when working with a desktop computer or a laptop, most computational servers and cloud platforms give the option or require the user to choose the memory allocated. Given it is common practice to over-allocate memory out of caution, we investigated the impact of memory allocation on carbon footprint in bioinformatics (**Figure 3, Supplementary table 1**).

**Figure 3:**
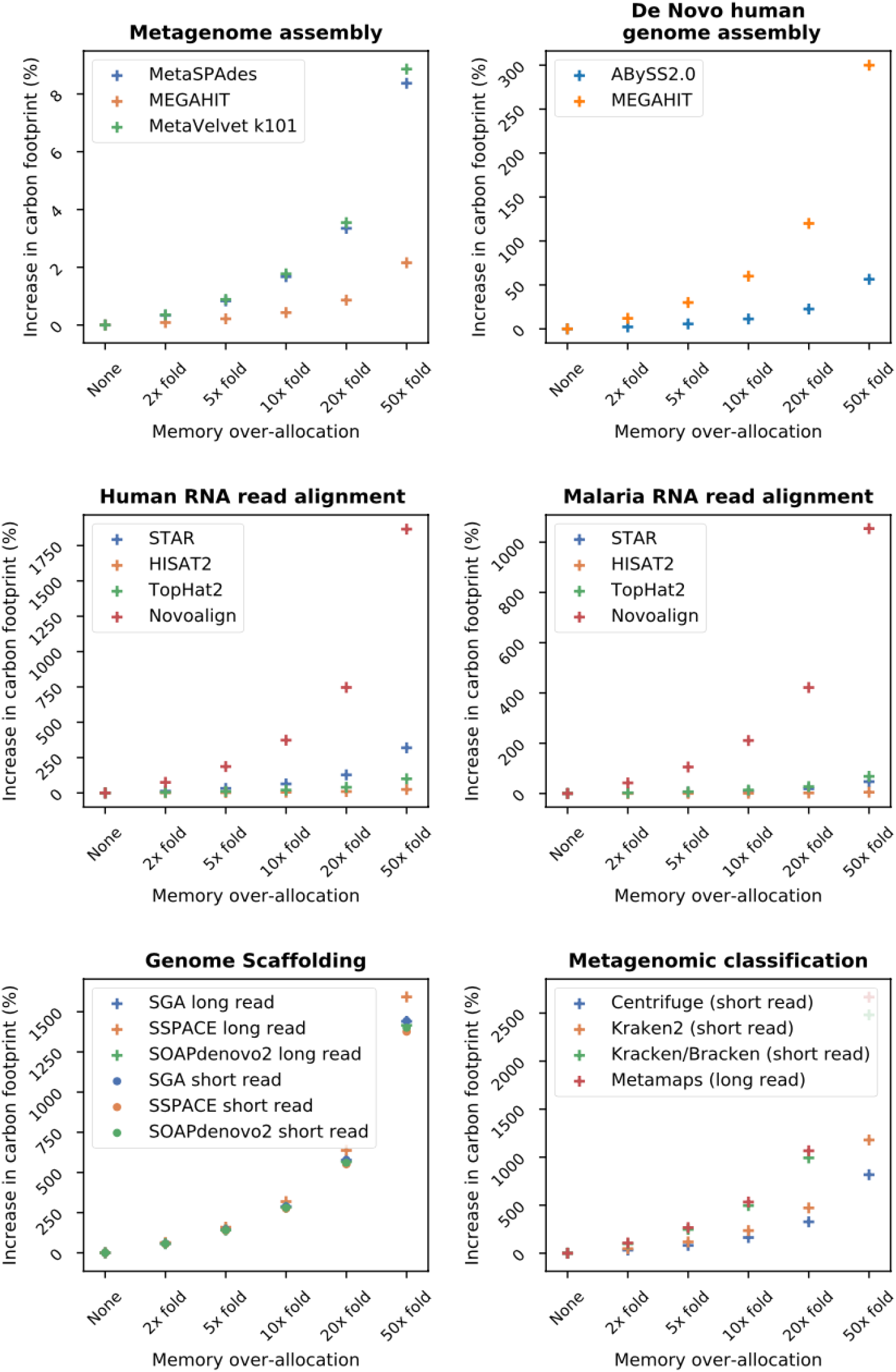
Over-allocating memory increases a given algorithm’s carbon footprint. Each plot details the percentage increase in carbon footprint as a function of memory overestimation for a variety of bioinformatic tools and tasks. The numerical data is available in **Supplementary Table 1**.

We showed that, while increasing the allocated memory always increases the carbon footprint, the effect is particularly significant for tasks with large memory requirements (**Figure 3, Supplementary table 1**). For example, in *de novo* human genome assembly, MEGAHIT had higher memory requirements than ABySS (6% vs 1% of total energy consumption); as a result, a five-fold over-allocation of memory increases carbon footprint by 30% for MEGAHIT and 6% for ABySS. Similarly, in human RNA read alignment (**Figure 3**), Novoalign had the highest memory requirements (37% of its total energy vs less than 7% for STAR, HISAT2, and TopHat2) and a 5x over-allocation in memory would increase its footprint by 186% compared to 32% for STAR, 2% for HISAT2, and 10% for TopHat2.

### Processors

We estimated the carbon footprint of a number of algorithms executed on both GPUs and CPUs. For cis-eQTL mapping (**Genome-wide association analysis**), we estimated that, compared to CPU-based FastQTL and LIMIX, using a GPU-based software like TensorQTL can reduce the carbon footprint by 96% and 99% and the running time by 99.63% and 99.99%, respectively (**Table 2**). For the codon modelling benchmark (**Phylogenetics**), utilising GPUs had a speedup factor of 93x and 13x when compared to 1 and 12 CPU cores, resulting in a decrease in carbon footprint of 75% and 84% respectively. These estimations demonstrate that GPUs can be well suited to both reducing running time and carbon footprint for algorithms.

However, there are situations where the use of GPUs can increase carbon footprint. Using a GPU for phylogenetic nucleotide modelling (**Phylogenetics**), instead of 8 CPU cores, decreased running time by 31% but also doubled the carbon footprint. We estimated that a single GPU would need to run the model in under four minutes in order to have the lowest carbon footprint for this analysis, as opposed to the 16 minutes it currently takes. Similarly, using a GPU for the phylogeographic modelling of the Ebola virus dataset (**Phylogenetics**) reduced the running time by 83% (6x speedup) when compared to the method with the lowest footprint (2 CPU cores) however, this increased carbon footprint by 84%. There are equations used for this estimation (**Supplementary Note 1**); however, a fast approximation can be used by scaling the running time of the GPU by the ratio of the power draw of the CPU cores to the GPU. For example, we compared the popular Xeon E5-2683 CPU (using all 16 cores) to the Tesla V100 GPU and found that, to have the same carbon footprint with both configurations, an algorithm needs to run 2.5 times faster on GPU than CPU.

## Discussion

We estimated the carbon footprint of various bioinformatic algorithms. Additionally, we investigated how memory over-allocation, processor choice and parallelisation affect carbon footprints, and showed the impact of transferring computations to hyperscale data centres.

This study made a series of important findings:

1. Limiting parallelisation can reduce carbon footprints. Especially when the running time reduction is marginal, the carbon cost of parallelisation should be closely examined.
2. Despite being often faster, GPUs don’t necessarily have a smaller carbon footprint than CPUs, and it is useful to assess whether the running time reduction is large enough to offset the additional power consumption.
3. Using currently optimised data centres, either local or cloud-based, can reduce carbon footprints by ~34% on average.
4. Substantial reductions in carbon footprint can be made by performing computations in energy-efficient countries with low carbon intensity.
5. Carbon offsetting, which consists of supporting GHG-reducing projects can be a way to balance the greenhouse gas emissions of computations. Although a number of cloud providers take part in this, [69], [72], [73], the real impact of carbon offsetting is debated and reducing the amount of GHG emitted in the first place should be prioritised.
6. Over-allocating memory resources can unnecessarily, and significantly, increase the carbon footprint of a task, particularly if this task has high memory usage already. To decrease energy waste, one should only allocate as closely as possible the required memory for a given job. Additionally, softwares could be optimised to minimise memory requirements, potentially moving some aspects to disk where energy usage is far lower.
7. A simple way to reduce the carbon footprint of a given algorithm is to use the most up to date software. We showed that updating common GWAS software reduced carbon footprint by 73%, indicating that this may be the quickest, easiest, and potentially most impactful way to reduce one’s carbon footprint.

There are a number of assumptions made when estimating the energy and carbon footprint of a given computational algorithm. These assumptions, and the associated limitations, have been discussed in detail within Lannelongue et al. [70]. A particularly important limitation of our study is that many of the carbon footprints estimated are from a single run of any given tool; however, many analyses have parameters that must be fine-tuned through trial and error, frequently extensively so. For example, in machine learning, thousands of optimisation runs may be required. We would stress that the total carbon footprint of a given project will likely scale linearly with the number of times each analysis is tuned or repeated, so a caveat to our estimations and the underlying published benchmarks is that the real carbon footprints could be orders of magnitude greater than that reported here.

Finally, the parameters needed to estimate the carbon footprint are often missing from published articles, such as running time, hardware information, and often software versions. If we are to fully understand the carbon footprint of the field of bioinformatics or computational research as a whole, there is a need for reporting this information as well as, ideally, for authors to estimate their carbon footprint using freely available tools.

## Conclusion

This study is, to the best of our knowledge, the first to estimate the carbon footprint for common bioinformatics tools. We further investigated how parallelisation, memory over-allocation, and hardware choices affect carbon footprints. We also show that carbon footprints could be reduced by utilising efficient computing facilities. Finally, we outline a number of ways bioinformaticians may reduce their carbon footprint.

## Methods

### Selection of bioinformatic tools

We estimated the carbon footprint of a range of tasks across the field of bioinformatics: genome and metagenome assembly, long and short reads metagenomic classification, RNA-seq and phylogenetic analyses, GWAS, eQTL mapping algorithms, molecular simulations and molecular docking algorithms (**Table 1**). For each task, we curated the published literature to identify peer-reviewed studies which computationally benchmarked popular tools. For our analysis, we used 10 published scientific papers. To be selected, publications had to report at least the running time and preferably the following: memory usage, and hardware used for the experiments, in particular the model and number of processing cores. We selected 10 publications for this study (**Table 1**). Besides, as we could not find suitable benchmarks to estimate the carbon footprint of cohort-scale eQTL mapping and RNA-seq quality control pipelines, we estimated the carbon footprint of these tasks using in-house computations. These computations were run on the Baker Heart and Diabetes Institute computing cluster (Intel Xeon E5-2683 v4 CPUs and a Tesla T4 GPU) and the University of Cambridge’s CSD3 computing cluster (Tesla P100 PCIe GPUs and Xeon Gold 6142 CPUs).

### Estimating the carbon footprint

The carbon footprint of a given tool was calculated using the framework described in Lannelongue et al. [70] and the corresponding online calculator www.green-algorithms.org. We present here an overview of the methodology.

Electricity production emits a variety of greenhouse gases, each with a different impact on climate change. To summarise this, the carbon footprint is measured in kilograms of CO_2_-equivalent (CO_2_e), which is the amount of carbon dioxide with an equivalent global warming impact as a mix of GHGs. This indicator depends on two factors: the energy needed to run the algorithm, and the global warming impact of producing such energy, called carbon intensity. This can be summarised by:

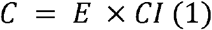

Where *C* is the carbon footprint (in kilograms of CO_2_e - kgCO_2_e), E is the energy needed (in W) and *CI* is the carbon intensity (in kgCO_2_e/W).

The energy needs of an algorithm are measured based on running time, processing cores used, memory deployed and efficiency of the data centre:

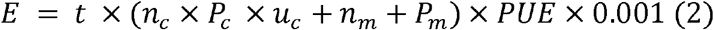

Where *t* is the run time (h), *n_c_* is the number of computing cores, used at *u_C_*%, the core usage factor (between 0 and 1), and each drawing a power *P_c_* (W). *n_m_* is the size of memory available (GB), drawing a power *P_m_* (W/GB). PUE is the Power Usage Effectiveness of the data centre.

The power drawn by a processor (CPU or GPU) is estimated by its Thermal Design Power (TDP) per core, which is provided by the manufacturer, and then scaled by the core usage factor *u_C_*. The power draw from memory was estimated to be 0.3725 W/GB [70]. The PUE represents how much extra energy is needed to run the computing facilities, mainly for cooling and lighting.

The carbon intensity (*CI*) varies between countries because of the heterogeneity in energy production methods, from 0.012 kgCO_2_e/kWh in Switzerland to 0.88 kgCO_2_e/kWh in Australia [74]. In order to be location-agnostic in this study, we used the global average value (0.475 kgCO_2_e/kWh [14]), unless otherwise specified.

### Reference values for carbon footprints

A quantity of carbon dioxide is not a metric most scientists are familiar with. To put the results presented here into perspective, we compare them to the impact of familiar activities. The first metric is the “tree-month”, defined as the number of months an average mature tree would take to fully sequester (absorb) an amount of carbon dioxide. A tree-month is defined as 0.917 kgCO_2_e [70]. Another way to contextualise a carbon footprint is to compare it with driving an average European car, which emits 0.175 kgCO_2_e/km [75], [76].

## Acknowledgement

We thank Kim van Daalen for the fruitful discussions about the impact of climate change on human health. JG was supported by a La Trobe University Postgraduate Research Scholarship jointly funded by the Baker Heart and Diabetes Institute and a La Trobe University Full-Fee Research Scholarship. LL was supported by the University of Cambridge MRC DTP (MR/S502443/1). This work was supported by core funding from: the UK Medical Research Council (MR/L003120/1), the British Heart Foundation (RG/13/13/30194; RG/18/13/33946) and the National Institute for Health Research [Cambridge Biomedical Research Centre at the Cambridge University Hospitals NHS Foundation Trust] [*]. This work was also supported by Health Data Research UK, which is funded by the UK Medical Research Council, Engineering and Physical Sciences Research Council, Economic and Social Research Council, Department of Health and Social Care (England), Chief Scientist Office of the Scottish Government Health and Social Care Directorates, Health and Social Care Research and Development Division (Welsh Government), Public Health Agency (Northern Ireland), British Heart Foundation and Wellcome. MI was supported by the Munz Chair of Cardiovascular Prediction and Prevention. This study was supported by the Victorian Government’s Operational Infrastructure Support (OIS) program. *The views expressed are those of the authors and not necessarily those of the NHS, the NIHR or the Department of Health and Social Care. JM is currently an employee of Genomics PLC.

## Availability of data and materials

The datasets used to support the conclusions of this article are available in supplementary information Additional file 1. The calculator used to estimate the carbon footprint is available at https://green-algorithms.org/, the code is available at https://github.com/GreenAlgorithms/green-algorithms-tool and the method behind it is described in Lannelongue et al [70].

## Supplementary materials

**Supplementary table 1:**
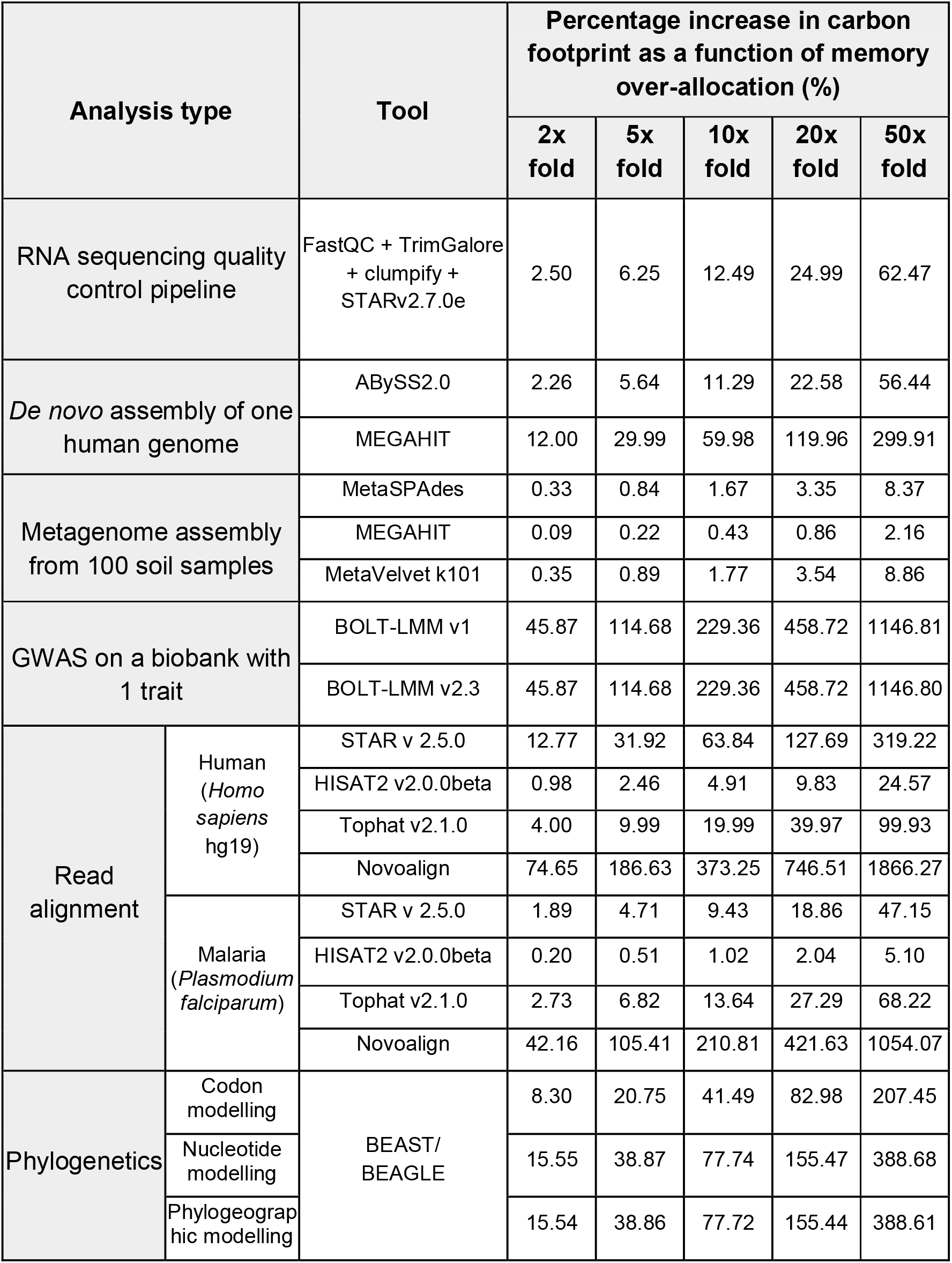
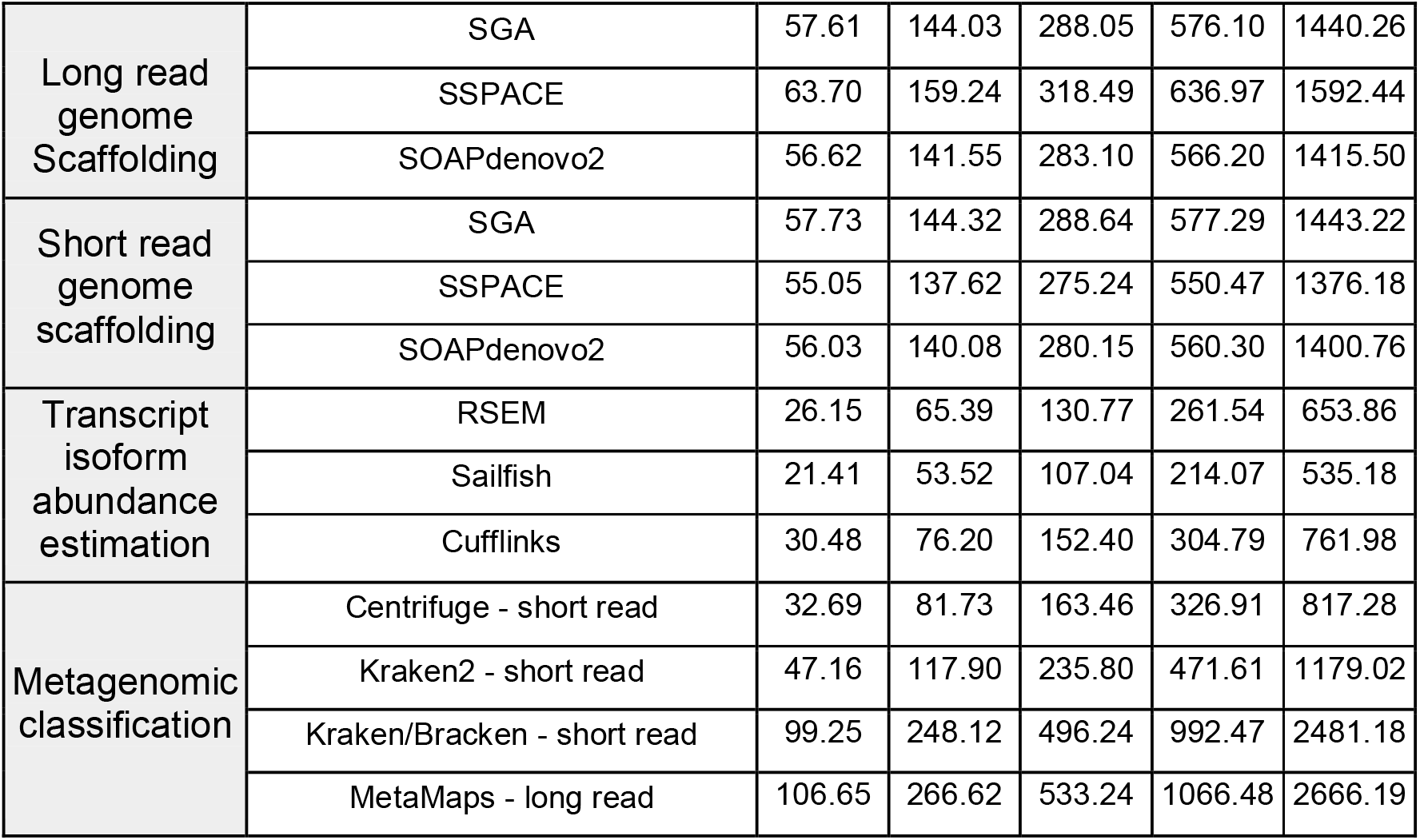
The percentage increase of carbon footprint as a function of memory over-allocation for a given algorithm.

| Analysis type |  | Tool | Percentage increase in carbon footprint as a function of memory over-allocation (%) |  |  |  |  |
| --- | --- | --- | --- | --- | --- | --- | --- |
|  |  |  | 2x fold | 5x fold | 10x fold | 20x fold | 50x fold |
| RNA sequencing quality control pipeline |  | FastQC + TrimGalore + clumpify + STARv2.7.0e | 2.50 | 6.25 | 12.49 | 24.99 | 62.47 |
| *De novo* assembly of one human genome |  | ABYSS2.0 | 2.26 | 5.64 | 11.29 | 22.58 | 56.44 |
|  |  | MEGAHIT | 12.00 | 29.99 | 59.98 | 119.96 | 299.91 |
| Metagenome assembly from 100 soil samples |  | MetaSPAdes | 0.33 | 0.84 | 1.67 | 3.35 | 8.37 |
|  |  | MEGAHIT | 0.09 | 0.22 | 0.43 | 0.86 | 2.16 |
|  |  | MetaVelvet k101 | 0.35 | 0.89 | 1.77 | 3.54 | 8.86 |
| GWAS on a biobank with 1 trait |  | BOLT-LMM v1 | 45.87 | 114.68 | 229.36 | 458.72 | 1146.81 |
|  |  | BOLT-LMM v2.3 | 45.87 | 114.68 | 229.36 | 458.72 | 1146.80 |
| Read alignment | Human (*Homo sapiens* hg19) | STAR v 2.5.0 | 12.77 | 31.92 | 63.84 | 127.69 | 319.22 |
|  |  | HISAT2 v2.0.0beta | 0.98 | 2.46 | 4.91 | 9.83 | 24.57 |
|  |  | Tophat v2.1.0 | 4.00 | 9.99 | 19.99 | 39.97 | 99.93 |
|  |  | Novoalign | 74.65 | 186.63 | 373.25 | 746.51 | 1866.27 |
|  | Malaria (*Plasmodium falciparum*) | STAR v 2.5.0 | 1.89 | 4.71 | 9.43 | 18.86 | 47.15 |
|  |  | HISAT2 v2.0.0beta | 0.20 | 0.51 | 1.02 | 2.04 | 5.10 |
|  |  | Tophat v2.1.0 | 2.73 | 6.82 | 13.64 | 27.29 | 68.22 |
|  |  | Novoalign | 42.16 | 105.41 | 210.81 | 421.63 | 1054.07 |
| Phylogenetics | Codon modelling | BEAST/ BEAGLE | 8.30 | 20.75 | 41.49 | 82.98 | 207.45 |
|  | Nucleotide modelling |  | 15.55 | 38.87 | 77.74 | 155.47 | 388.68 |
|  | Phylogeographic modelling |  | 15.54 | 38.86 | 77.72 | 155.44 | 388.61 |

|  |  |  |  |  |  |  |
| --- | --- | --- | --- | --- | --- | --- |
| Long read<br>genome<br>Scaffolding | SGA | 57.61 | 144.03 | 288.05 | 576.10 | 1440.26 |
|  | SSPACE | 63.70 | 159.24 | 318.49 | 636.97 | 1592.44 |
|  | SOAPdenovo2 | 56.62 | 141.55 | 283.10 | 566.20 | 1415.50 |
| Short read<br>genome<br>scaffolding | SGA | 57.73 | 144.32 | 288.64 | 577.29 | 1443.22 |
|  | SSPACE | 55.05 | 137.62 | 275.24 | 550.47 | 1376.18 |
|  | SOAPdenovo2 | 56.03 | 140.08 | 280.15 | 560.30 | 1400.76 |
| Transcript<br>isoform<br>abundance<br>estimation | RSEM | 26.15 | 65.39 | 130.77 | 261.54 | 653.86 |
|  | Sailfish | 21.41 | 53.52 | 107.04 | 214.07 | 535.18 |
|  | Cufflinks | 30.48 | 76.20 | 152.40 | 304.79 | 761.98 |
| Metagenomic<br>classification | Centrifuge - short read | 32.69 | 81.73 | 163.46 | 326.91 | 817.28 |
|  | Kraken2 - short read | 47.16 | 117.90 | 235.80 | 471.61 | 1179.02 |
|  | Kraken/Bracken - short read | 99.25 | 248.12 | 496.24 | 992.47 | 2481.18 |
|  | MetaMaps - long read | 106.65 | 266.62 | 533.24 | 1066.48 | 2666.19 |

**Supplementary table 2:** The carbon footprint of hardware changes and parallelisation, using benchmarks from Beale et al [32].

| Task | Algorithm | Number of CPU cores or GPU devices | Running time (hours) | Carbon footprint (kgCO<sub>2</sub>e) |
| --- | --- | --- | --- | --- |
| Codon substitution modelling | BEAST/<br>BEAGLE | 1 | 7.75 | 0.066 |
|  |  | 2 | 4.17 | 0.069 |
|  |  | 4 | 2.42 | 0.078 |
|  |  | 6 | 1.72 | 0.083 |
|  |  | 8 | 1.42 | 0.091 |
|  |  | 10 | 1.25 | 0.10 |
|  |  | 12 | 1.08 | 0.10 |
|  |  | 1 GPU<br>2 GPU | 0.08<br>0.06 | 0.017<br>0.023 |
| Nucleotide substitution modelling | BEAST/<br>BEAGLE | 2 | 0.67 | 0.012 |
|  |  | 4 | 0.43 | 0.015 |
|  |  | 6 | 0.40 | 0.020 |
|  |  | 8 | 0.39 | 0.026 |
|  |  | 10 | 0.43 | 0.035 |
|  |  | 12 | 0.43 | 0.042 |
|  |  | 1 GPU<br>2 GPU | 0.27<br>0.19 | 0.054<br>0.076 |
| Phylogeographic modelling | BEAST/<br>BEAGLE | 2 | 3.86 | 0.070 |
|  |  | 4 | 3.73 | 0.13 |
|  |  | 6 | 3.69 | 0.18 |
|  |  | 8 | 3.71 | 0.24 |
|  |  | 10 | 3.68 | 0.30 |
|  |  | 1 GPU<br>2 GPU | 0.64<br>0.54 | 0.13<br>0.22 |

### Supplementary Note 1

#### Estimating the running time at which a GPU has a lower carbon footprint

From rearranging the Green Algorithms carbon footprint formula it can be shown that the running time at which GPU has a lower carbon footprint is:

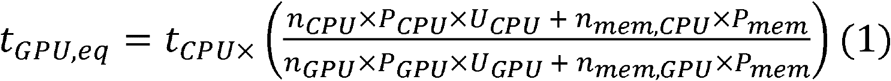

Where, *n_CPU_* is the number of CPU cores, *n_GPU_* is the number of GPUs, *P_CPU_* is the power drawn by the CPU cores. *P_GPU_* is the power drawn by the GPU. *U_CPU_* is the core usage factor for the CPU. *U_GPU_* is the usage factor of the GPU. *n_mem,CPU_* is the amount of memory (GB) utilised when running the CPU, *n_mem,GPU_* is the amount of memory (GB) utilised when running the GPU. *P_mem_* is the power draw for memory. *t_GPU,eq_* is the running time when the GPU would have the same carbon footprint as the CPU, and *t_CPU_* is the running time of the CPU. If the GPU implementation is to have a lower carbon footprint, it must finish within the time *t_GPU,eq_*.

When ignoring memory and utilising 1 CPU and 1 GPU with identical core usage factors, this simplifies to:

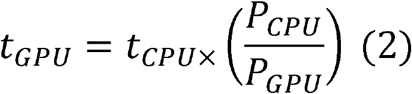

Where, *t_CPU_* is scaled by the ratio of the power required to utilise the CPU to the GPU.

#### Descriptions of additional files

**Additional file 1:** Hardware details for each analysis presented in this manuscript.

**Additional file 2:** The ratio of RNA reads per million and ratio of CPU time of 10 random in house PBMC samples, from the RNA sequencing quality control pipeline task.

## Notes

### Competing Interest Statement

The authors have declared no competing interest.

